# Inferring Cognitive State Underlying Conflict Choices in Verbal Stroop Task Using Heterogeneous Input Discriminative-Generative Decoder Model

**DOI:** 10.1101/2022.11.28.518256

**Authors:** Mohammad R. Rezaei, Haseul Jeoung, Ayda Gharamani, Utpal Saha, Venkat Bhat, Milos R. Popovic, Ali Yousefi, Robert Chen, Milad Lankarany

**Affiliations:** Department of Biomedical Engineering, University of Toronto, ON, Canada; Krembil Research Institute-University Health Network (UHN) Toronto, ON, Canada; KITE Research Institute-University Health Network (UHN), ON, Canada; School of Medicine, Stanford University, CA, USA; Worcester Polytechnic Institute, MA, USA; Department of Psychiatry, University Health Network University of Toronto, Toronto, Canada

**Keywords:** Cognitive state decoding, State-space model, Subthalamic nucleus, Verbal Stroop task, Parkinson’s disease

## Abstract

The subthalamic nucleus (STN) of the basal ganglia interacts with the medial prefrontal cortex (mPFC) and shapes a control loop, specifically when the brain receives contradictory information from either different sensory systems or conflicting information from sensory inputs and prior knowledge that developed in the brain. Experimental studies demonstrated that significant increases in theta activities (2-8 Hz) in both the STN and mPFC as well as increased phase synchronization between mPFC and STN are prominent features of conflict processing. While these neural features reflect the importance of STN-mPFC circuitry in conflict processing, a low-dimensional representation of the mPFC-STN interaction referred to as a cognitive state, that links neural activities generated by these sub-regions to behavioral signals (e.g., the response time), remains to be identified. Here, we propose a new model, namely, the heterogeneous input discriminative-generative decoder (HI-DGD) model, to infer a cognitive state underlying decision-making based on neural activities (STN and mPFC) and behavioral signals (individuals’ response time) recorded in 10 Parkinson’s disease patients while they performed a Stroop task. PD patients may have conflict processing which is quantitatively (may be qualitative in some) different from healthy population. Using extensive synthetic and experimental data, we showed that the HI-DGD model can diffuse information from neural- and behavioral data simultaneously and estimate cognitive states underlying conflict and nonconflict trials significantly better than traditional methods. Additionally, the HI-DGD model identified which neural features made significant contributions to conflict and non-conflict choices. Interestingly, the estimated features match well with those reported in experimental studies. Finally, we highlight the capability of the HI-DGD model in estimating a cognitive state from a single trial of observation, which makes it appropriate to be utilized in closed-loop neuromodulation systems.

**Highlights:**

- Research highlight 1
- Research highlight 2

## 1. Introduction

The subthalamic nucleus (STN) of the basal ganglia (BG) and prefrontal cortex (PFC) are essential brain structures in the decision-making processes Bonnevie and Zaghloul (2019); Drummond and Chen (2020). The interaction between the STN and the PFC results in slower and more controllable decisions during conflict tasks which has been previously demonstrated Drummond and Chen (2020); Bočková et al. (2011); Bonnevie and Zaghloul (2019); Frank (2006). The presence of ambiguity in selection choices or conflict choices, activates the cortex which sends excitatory inputs to the STN via the hyper-direct pathway Frank (2006); Haynes and Haber (2013); Aron et al. (2016); Kelley et al. (2018); Bočková et al. (2020). In response, the STN sends excitatory inputs to the globus pallidus internus (GPi). GPi sends inhibitory inputs to the thalamus to suppress subsequent motor actions that lead to wrong decisions Wessel et al. (2019); Frank (2006); Aron et al. (2016); Kelley et al. (2018). This inhibition acts as a brake that slows the decision-making process down and helps to prevent impulsive decisions Frank (2006); Aron et al. (2016); Kelley et al. (2018); Brittain et al. (2012).

Despite this mechanistic understanding of the role of the PFC-STN neuronal circuit in decision-making with competing options, a low-dimensional representation of the PFC-STN interactions that links neural activities generated by these sub-regions to behavioral signals (e.g., the response time) remains to be identified. This cognitive state can reveal the state of the brain during decision-making. Such a low dimensional representation, also known as the latent brain state, was identified in other situations, e.g., the mood state in patients with major depression Sani et al. (2018), the state underlying cognitive deficits Basu et al. (2021), the state of speech intention Rezaei et al. (2022a), the learning state in memory Suzuki and Brown (2005), the state of attention in eye-metrics Marshall (2007), and the state underlying pain Villemure and Bushnell (2002). In this work, we propose a probabilistic model that accounts for the dynamics of the cognitive states underlying conflict processing in the verbal Stroop task. Additionally, by developing an inference method, we estimate this cognitive state that links temporal changes in the neural activities observed in local field potentials recordings from the STN and EEG recordings of PFC to the behavioral signal measured by participants’ response time.

Various methods including state-space models (SSMs) Nolte et al. (2004); Truccolo et al. (2005); Rezaei et al. (2021) and machine learning approaches Güçlütürk et al. (2017); Rezaei et al. (2018); ? have been used to model cognitive states using neural or behavioral data Yousefi et al. (2019); Basu et al. (2021); Sani et al. (2018); Prerau et al. (2008); Coleman et al. (2011). An important key assumption behind these approaches is that dynamics in the latent brain states, hereafter known as cognitive states, are encoded in a subset of features of the neural activity and behavioral signals. Therefore, it can be decoded from either neural recordings or behavioral signals.

Despite the usefulness of these methods in decoding cognitive states, they suffer from several limitations. For high-dimensional neural data (recorded by several electrodes over time), these models become impractical and computationally inefficient for building accurate receptive-field models for individual neurons or sources Goodwin et al. (2001); Van Der Merwe (2004); Abellán-Nebot et al. (2009); Robert et al. (2010). Building accurate receptive-field models requires additional modeling steps, such as spike sorting or clusterless models, which add complexity and computational expense to the process Wood et al. (2004). Moreover, individual neural receptive-field models do not contain information about the co-activity across neurons that encapsulate essential information about interactions among different brain regions of the brain Panzeri et al. (2015); Ruff et al. (2018). For example, traditional models often ignore the coherence and synchrony between different channels of neural activity for decoding a cognitive state Basu et al. (2021).

In addition, traditional models do not use all possible sources of information at the same time for decoding a cognitive state Sani et al. (2018); Yousefi et al. (2019); Basu et al. (2021); Sani et al. (2018, 2021). The existing methods utilize behavioral signals as a noisy observation of the designated cognitive state. Then, the cognitive state is solely decoded from neural signals. For example, Yousefi, et al Yousefi et al. (2019) considered the response time of participants performing a multisource interference task (MSIT) as a noisy measure of cognitive flexibility state and inferred it from LFP signals in a separate step Bush and Shin (2006). Therefore, this strategy of decoding a cognitive state employs only one source of information and may lead to suboptimal results.

In addition to the above limitations, one should notice that the conventional data analysis methods used in experimental neuroscience cannot decode cognitive states underlying neural and behavioral data. These methods detect neural features which are significantly different in distinct cognitive tasks, e.g., the power of theta band (in both STN and PFC) in conflict and nonconflict tasks Ghahremani et al. (2018); Zeng et al. (2021). Such traditional analyses are performed based on a group average from different experimental trials of an experiment, making these methods impractical for single-trial analysis.

To address above mentioned limitations, we propose a Bayesian-based model, the heterogeneous input discriminative-generative decoder (HI-DGD), which can combine information from multiple sources with different sampling rates. The HI-DGD consists of

1. A state transition model that controls the cognitive state evolvement over time.
2. A generative model that represents the behavioral signal as a function of the cognitive state.
3. A discriminative model that formulates the cognitive state as a function of (high-dimensional) neural activity features.

This hybrid approach will allow us to simultaneously combine simple generative models of behavior as a function of the underlying cognitive state with information encoded in complex patterns of high-dimensional neural activity. By using HI-DGD, we show that cognitive states underlying conflict and non-conflict tasks can be inferred from individual trials of the experiment accurately. Moreover, calculated model parameters reveals significant features of neural activities that classify conflict and non-conflict choices. Interestingly, these features have been proven experimentally, demonstrating the fidelity of our approach.

## 2. Heterogeneous Input Discriminative-Generative Decoder (HI-DGD) model

Here we propose the heterogeneous input discriminative-generative decoder (HI-DGD) model. HI-DGD model infers a cognitive state, ***x***_*k*_, that is a low-dimensional representation of conflict processing state underlying observed neural activity, ***s***_1:*k*_, and behavioral signals, ***z***_1:*k*_. A graphical representation of the HI-DGD model is shown in Figure1.B. The HI-DGD model uses three processes to infer a cognitive state that links neural activities and behavioral observations:

1. A state transition process that represents the evolution of the states in time, which follows a random walk model Gerstein and Mandelbrot (1964), *p*(***x***_*k*_|***x***_*k*–1_).
2. A discriminative process to link neural activity and the cognitive state as a function of high-dimensional observations, *P*(***x***_*k*_|***s***_*k*_).
3. A generative process that links the cognitive state to behavioral signals, *P*(***z***_*k*_|***x***_*k*_).

**Figure 1:**
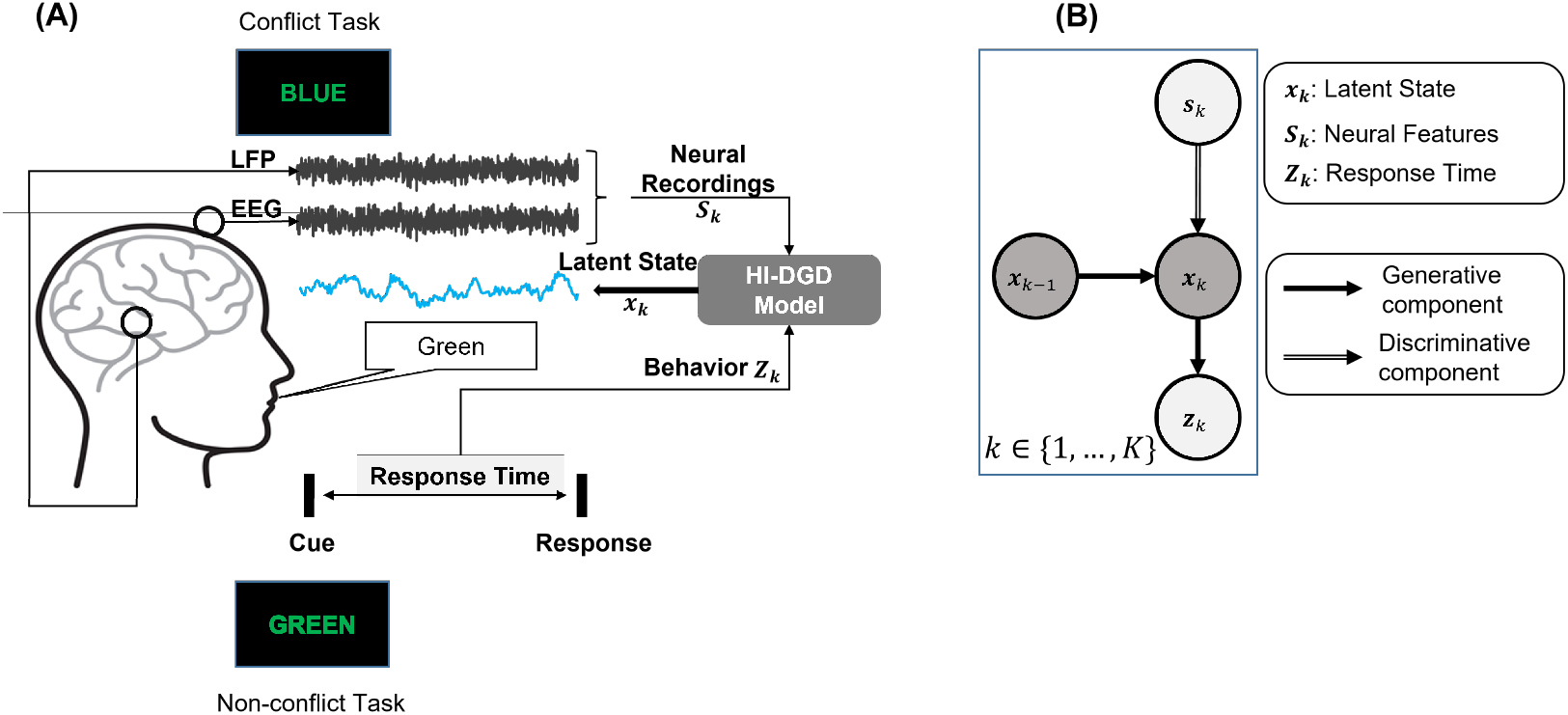
HI-DGD encodes the cognitive state associated with conflict processing from neural activities of STN and mPFC and the patient response time. A) A schematic view of the encoding mechanism. Each trial of the experiment begins with a cue signal, after which a color word appears on the screen. The subject should respond with the ink color of the words and ignore their meaning. As illustrated, the subject would speak (RED) whether the word is (RED) or (GREEN). A one-to-one ratio of conflict and non-conflict trials was used in this study. B) The graphical view of the HI-DGD model. ***s**_k_, **z**_k_, **x**_k_* represent the neural features, behavioral features, and the inferred cognitive state at time *k*; respectively.

Assume that ***s**_k_* represents high dimensional observations, at time index *k*, and ***z**_k_* represents all other observation elements which are usually low dimensional or very sparse Suzuki and Brown (2005); Truccolo et al. (2005). Given ***s**_k_* and ***z**_k_*, our goal for the HI-DGD model is to infer the posterior distribution over cognitive states, ***x**_k_*, as written below

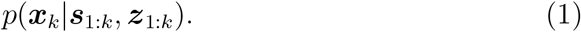

This equation defines the conditional density function of the current value of the cognitive state given all sources of observations from *k* = 1. Following Rezaei et al. (2022b), equation 1 can be approximated as follow

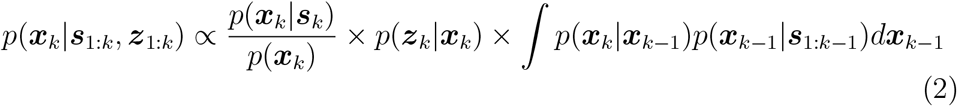

where the nominator of the first term is called the prediction process. The second term of equation 2, *p*(***z**_k_*|***x**_k_*), is the conditional density of the behavior observation, i.e., response time, given the current value of the state. The integral term of equation 2 is the one-step prediction density by the posterior function from the last time-step derived by the Chapman-Kolmogorov equation Karush (1961).

The state transition process is defined by a set of free parameters ***β*** as

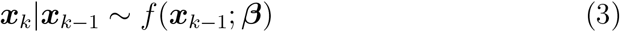

The observation processes are defined by

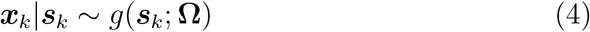

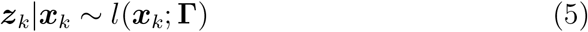

where equations 4 and 5 indicate observations processes for the neural and the behavioral signals, respectively.

Most of the previous studies in cognitive state modeling Sani et al. (2018); Basu et al. (2021); Yousefi et al. (2019); Sani et al. (2021) considered a two-step process, namely, encoding and decoding, to identify the relationship between brain dynamics and behavior through a latent dynamical variable. In contrast to these works, the HI-DGD model infers the underlying cognitive state using both neural- and behavioral-observations simultaneously. In this way, the HI-DGD can use information from both behavior and neural data simultaneously to estimate cognitive dynamics. In addition, the HI-DGD model can combine information at different temporal scales through the process defined in equations 3-5 to infer the underlying cognitive state. This property will make the solution flexible and powerful in decoding a cognitive state from different sources of information where each source carries information in different temporal scales.

### 2.1. Marked-point process observation model for modeling behavioral signals

In the present work, we utilized a marked point-process (MPP) observation model Deng et al. (2015) for modeling the behavioral signals, both response times and trial types (conflict or non-conflict) in the Stroop task Scarpina and Tagini (2017). Each participant’s response time is considered an event. We define *r_k_* ∈ {0,1} as the random variable representing the onset of the participant’s response time. At each time step *k, r_k_* = 1 indicates the onset of the participant’s response and *r_k_* = 0 otherwise. Additionally, to differentiate between conflict and non-conflict tasks, we define another random variable, referred to as a mark, *m_k_* ∈ {-1,1}, in the MPP theory Deng et al. (2015). The variable *m_k_* can take two values, 1 and −1, where 1 represents a conflict decision, and −1 indicates a non-conflict choice. The distribution of the *r_k_* and *m_k_* represents the MPP observation process associated with the behavioral signals.

For the MPP observation, the joint mark intensity function of the process is defined by

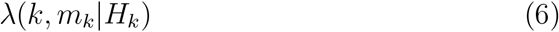

where λ defines the instantaneous probability of observing an event at time index *k* with mark *m_k_* given the full history of events up to time *k, H_k_*. Using the factorization rule, the joint distribution of the mark values and events is proportional to

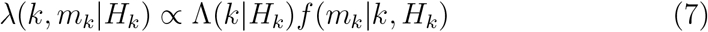

where Λ(*k*|*H_k_*) = ∑_*m*_ λ(*k,m_k_* | *H_k_*) is the conditional intensity function (CIF) for the underlying event process and *f* (*m_k_*|*k,H_k_*) is the conditional intensity of the marks for specific events. Hereafter Λ(*k*|*H_k_*) is written as Λ_*k*_ for the sake of simplicity. In the case of observation processes characterized by the response time accompanied by a two choices decision like conflict and non-conflict choices, we can define the following distribution

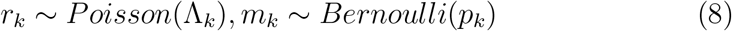

Note that there are only two types of tasks, therefore *m_k_* can be fully represented by a Bernoulli random variable. To be consistent with the general linear model (GLM) theory, we used the ”log” and ”logit” link functions to describe the Poisson and Bernoulli distributions, respectively. Therefore, the predictor can be defined as a function of the cognitive state by

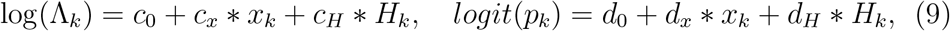

where {*c*_0_, *d*_0_, *c_x_, d_x_, c_H_, d_H_*} are the observation process free parameters Amidi et al. (2019) and *H_k_* is the history term of previous events. Therefore, the observation likelihood can be formulated as

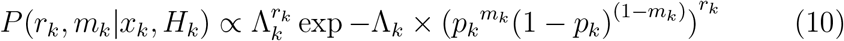

The first term, 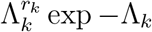, characterizes the distribution of observing an event (the speech onset) and the remaining term, (*p_k_^mk^*(1 – *p_k_*)^(1–*m_k_*))r_k_^, characterizes the distribution of mark values (trial type) given an event time. The remaining continuous observations such as LFP and EEG neural features during the cognitive task can be modeled as Gaussian observation processes (unless another type of observation is preferred) defined as

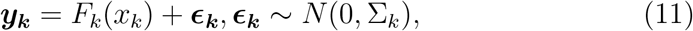

where *F_k_*(.) maps the state values to the observations and *ϵ_k_* is additive white Gaussian noise with covariance matrix ∑_*k*_. Using extensive synthetic data, we verified the performance of the HI-DGD model in inferring a latent state in Appendix B and FigureB.7.

### 2.2. Feature extraction and data processing

To reach a set of representative features relevant to conflict processing in the brain, we replicated the procedure proposed in Ghahremani et al. (2018). We wrote a customized Python script along with using the MNE library Gramfort et al. (2013) to analyze the LFP/EEG signals data. All LFP/EEG data were down-sampled to 1,000 Hz and filtered by a band-passed filter (between 1 and 400 Hz).

#### 2.2.1. Power spectrum

Following Ghahremani et al. (2018), we use time-frequency features from 3 seconds before to 2 seconds after the cue time. The LFP/EEG was decomposed into the time-frequency domain using a bank of band-pass filters for the 2-100Hz in steps of 1Hz. Then, the frequency features are normalized with the baseline power of 3 to 2 seconds before the cue time using

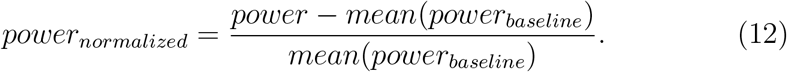

We quantified the power spectrum features for delta-theta (2–8 Hz), alpha-l (8–12 Hz), alpha-h (12–24 Hz), low-beta (24-34 Hz), gamma-l (30–55 Hz), and gamma-h (65–100 Hz) frequency bands by averaging power spectrum over each frequency band.

#### 2.2.2. Phase coherency

The STN and mPFC phase synchronization values (Coh) at (*k, f*), *Coh*(*k, f*), were estimated by projecting the phase difference between STN (LFP) and cortex (EEG) by phase coherency measure described in Lutz et al. (2002); Nolte et al. (2004) as follows.

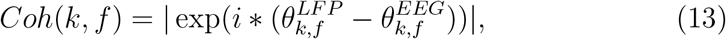

here *k* is the time step (here we considered 10ms time resolutions for the time steps) and *f* is the frequency. The range of *Coh*(*k, f*) is from 0 to 1, where 0 indicates random phase differences between STN and medial prefrontal cortex (mPFC), and 1 indicates identical phase differences.

#### 2.2.3. Behavioral signals

For behavioral recording, a microphone of an Apple MacBook recorded the patients’ voices. The speech onset time considered the response time and also measured the patient’s performance based on his/her ability in making the correct decision. Trials with response time > 2.0 seconds or < 0.3 seconds and those with noise or unclear/incorrect speech responses were excluded from the analysis used in Ghahremani et al. (2018).

## 3. Results

Previous studies demonstrated that conflict processing is associated with statistical changes in the power and the phase of low-frequency oscillations (2-8Hz - delta and theta bands) in the STN and medial prefrontal cortex (mPFC), as well as in behavior such as the participants’ response-time Ghahremani et al. (2018); Zeng et al. (2021). We analyzed LFPs, EEGs, and response time of 6 Parkinson’s disease (PD) patients who underwent DBS surgeries recruited in a verbal Stroop task. A schematic view of neural feature extraction from LFP signals recorded from STN and EEG signals recorded from mPFC is shown in Figure3. A. This figure includes the inferred cognitive state from neural and behavioral signals. Details regarding the patients and experimental procedures reported in a previous study that studied the same cohort of patients Ghahremani et al. (2018) (patients 3, 5-13 in Table 1 in Ghahremani et al. (2018)). In the present work, we selected 6 out of 10 patients to test the accuracy of our proposed algorithm for individual trials. The selection criterion was based on the accuracy of the baseline model; we excluded those patients with low accuracy (~ 50% accuracy) in distinguishing conflict and non-conflict trials by the baseline model discussed in section 3.4. Moreover, we compared the accuracy of the proposed HI-DGD model with that of the baseline model including all 10 patients. As shown in supplementary results - Figure C.8, the accuracy of the proposed model is significantly higher than that of the baseline model (p-value ≤ 0.001). To this end, we used neural and behavioral signals to find a low-dimensional cognitive state underlying conflict processing in the selected participants.

**Table 1:** Neural featured used in the HI-DGD model.

|  |  |  |  |
| --- | --- | --- | --- |
| F01 | Power STN 2-8Hz | F10 | Power mPFC 24-34Hz |
| F02 | Power STN 8-12Hz | F11 | Power mPFC 34-55Hz |
| F03 | Power STN 12-24Hz | F12 | Power mPFC 65-95Hz |
| F04 | Power STN 24-34Hz | F13 | Phase Coh. STN-mPFC 2-8Hz |
| F05 | Power STN 34-55Hz | F14 | Phase Coh. STN-mPFC 8-12Hz |
| F06 | Power STN 65-95Hz | F15 | Phase Coh. STN-mPFC 12-24Hz |
| F07 | Power mPFC 2-8Hz | F16 | Phase Coh. STN-mPFC 24-34Hz |
| F08 | Power mPFC 8-12Hz | F17 | Phase Coh. STN-mPFC 34-55Hz |
| F09 | Power mPFC 12-24Hz | F18 | Phase Coh. STN-mPFC 55-95Hz |

### 3.1. Identifying optimal kernel of neural features using behavioral signals

To identify how features of recorded neural activities are linked to the behavioral signal, we, first, utilized a Poisson probabilistic model that represents the response time as a function of neural features. This is a traditional probabilistic model that frequently has been used to identify the relationship between inputs and the time of an event (here, patient response time) Suzuki and Brown (2005); Truccolo et al. (2005). By considering *k_c_* as the start time of a trial, *ψ_k_* ∈ ℝ^*M*^ as the feature vector extracted from neural activities, and *M* as the number of the neural features for each time index, we define the response time *r_k_* at time index *k* with a Poisson probabilistic model as

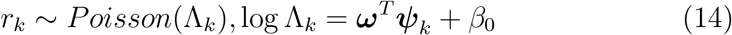

where the Λ_*k*_ is the conditional intensity function that links the neural features to the response time *r_k_* with a Poisson distribution. The parameter *β*_0_ governs the baseline response time, whereas ***ω*** represents the weight by which the subject response time is related to the neural features. The logarithmic relationship imposes an empirical limitation of human response time into the model Prerau et al. (2008, 2009). The model parameters {***ω***, *β*_0_} can be optimized by a generalized linear model (GLM) with a proper link function and additive noise Truccolo et al. (2005). Figure 2.B shows the absolute weight of the extracted neural features (**ω**) in representing the response times for the patients. As the time passes from the cue time, the importance of the neural features that represent the response time of a patient decreases, and the variability of the features increases. To identify the optimal kernel width during which neural features represent response times, we calculated the log-likelihood (LL) of the optimized model and plotted the measure for different windows in Figure2.C. The LL value represents the goodness of fit for the behavioral model. The higher the value of the LL, the better a model fits to a dataset. The LL converges to its maximal value after [400 – 500]ms of observing neural features after the cue time. This means that increasing the time window for neural features is not statistically significant for the model fitting. Therefore, the most relevant neural features that represent a participants’ response times, i.e. behavioral signal, lie within the 500ms time window after the cue time.

**Figure 2:**
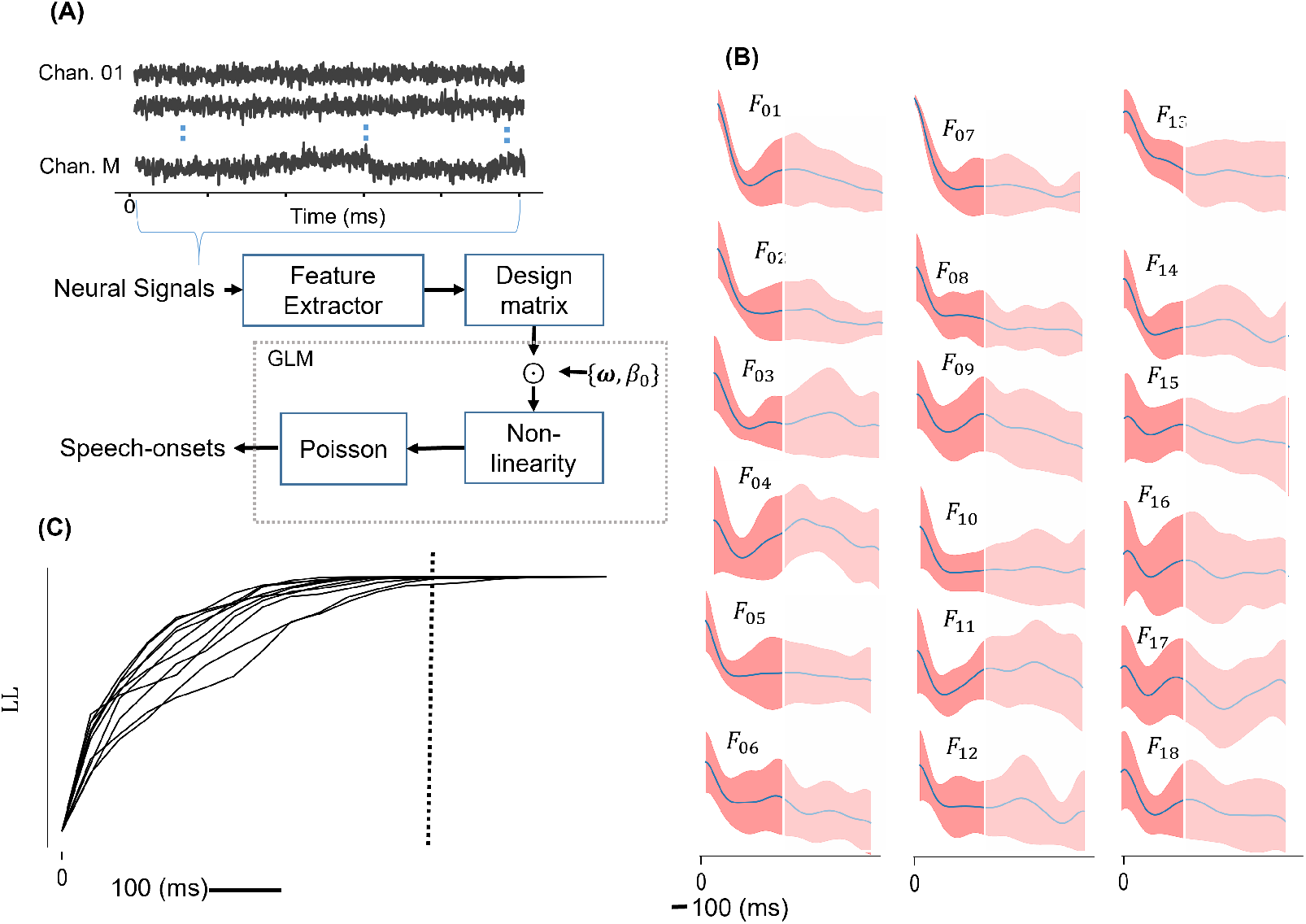
Behavioral model and optimal kernels. A) The schematic view of creating the design matrix for the lasso-GLM model. B) the mean relative importance of the extracted neural features (***ω***) in time to the response time. The neural features are defined in Table1 C) The log-likelihood (LL) of the lasso-GLM model vs the time window length of the neural features after the cue time. The LL of the model stops growing after [400 – 500]ms of observing neural features windows.

### 3.2. HI-DGD model parameters inference using neural activity and behavioral signal

Building on the analysis from the previous section and experimental evidence related to conflict processing representation in both neural and behavioral signals, we used the HI-DGD model to infer a cognitive state that represents the dynamics underlying conflict/non-conflict processing from neural and behavioral signals (response time). Figure1.b shows a graphical representation of the HI-DGD model. In HI-DGD, {*x_k_*, ***s***_*k*_, ***z***_*k*_} represents the cognitive state, neural features, and the behavioral features at time index k, respectively. The HI-DGD model uses ***s**_k_* and ***z**_k_* as noisy representations of the conflict/non-conflict processing dynamics and continuously infers a cognitive state *x_k_*.

Therefore, the objective for the HI-DGD model optimization is to find a parameter set *θ* = {*β*, Ω, Γ} that maximizes the joint distribution of observations, neural and behavioral signals, and the cognitive state, *P*(*x_k_*, *z_k_*, ***s**_k_*; ***θ***), for all time steps *K*. To meet this objective, as shown in Figure3.A, we optimized the HI-DGD model parameters by maximizing the following function

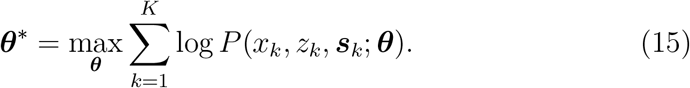

**Figure 3:**
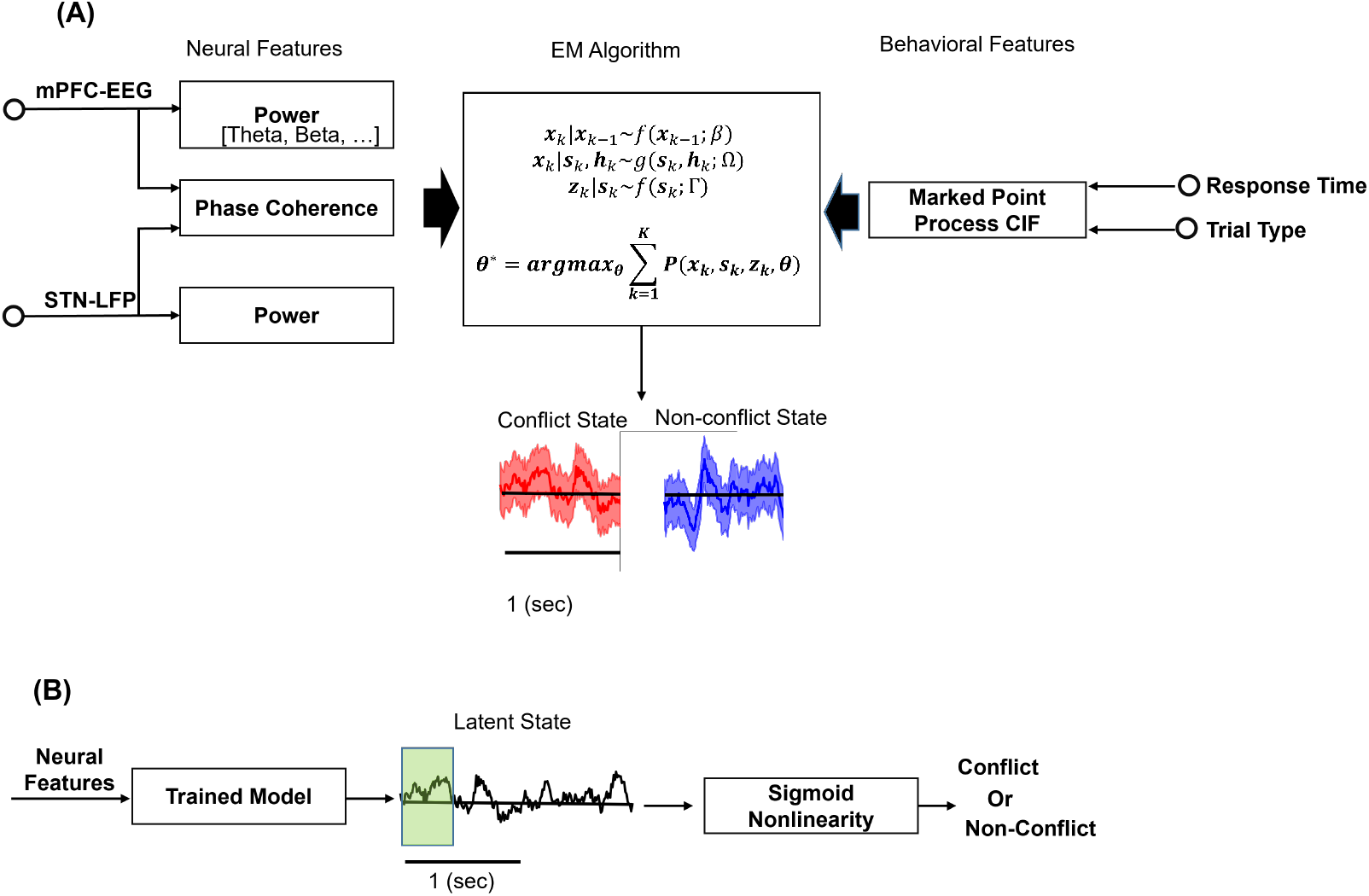
HI-DGD model identification and inference. A) A schematic representation of HI-DGD model identification. Neural feature extraction from LFP signals recorded from the STN and EEG signal recorded from the mPFC. We quantified the frequency band features by averaging them in frequency bands delta-theta (2–8 Hz), alpha (8–12 Hz), beta-l (12–24 Hz), beta-h (24–34 Hz), gamma-1 (34–55 Hz), and gamma-2 (65–95 Hz). We extracted power and phase coherency features according to what is described in the methods section. Both neural and behavioral features are passed to the HI-DGD model and the EM algorithm optimizes the identifies the optimal values for the model parameters set ***theta\****. The HI-DGD model output is an inferred cognitive state that represents the conflict processing in the brain. B) The conflict state prediction from the HI-DGD outputs (the inferred cognitive state). The statistics of the first 500 ms of the inferred cognitive state by the HI-DGD pass to a logistic regressor to predict the probability of the conflict and non-conflict states.

Since both model parameters and the true cognitive state are unknown, one cannot simply use a maximum likelihood approach for finding the optimal parameters. To address this issue, we used expectation-maximization (EM) to estimate the model parameters. The EM algorithm is an established method for maximum likelihood estimation of model parameters in the presence of a latent process or missing observations Wu et al. (2013). Other approaches including fully Bayesian or variational-Bayesian approaches, can be applied to our modeling approach in the case that a prior distribution for the model parameters can be defined Kingma and Welling (2013). In Appendix A. we derived the EM optimization for the HI-DGD model which recursively updates the model parameters ***θ***^(*r*+1)^ = {***β***^(*r*+1)^, **Ω**^(*r*+1)^, **Γ**^(*r*+1)^}, where superscript *r* indicates an iteration of the EM. Based on the updated posterior distribution of iteration *r*, the objective function *Q* is defined as

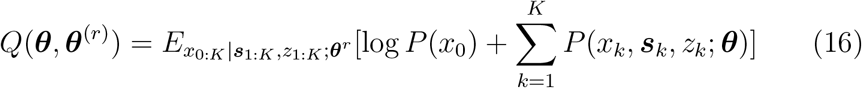

With an update rule as

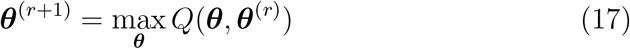

Further details of the function *Q* are provided in Appendix A. The value of the objective function for the EM algorithm is shown in Figure4.A., wherein the increasing trend in *Q* values shows that the new parameters set for the HI-DGD, ***θ***^*r*+1^, improves log-likelihood for the joint distribution of 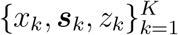. The *Q* values eventually reaches a steady state at around 15 iterations and gives the optimal model parameters sets.

**Figure 4:**
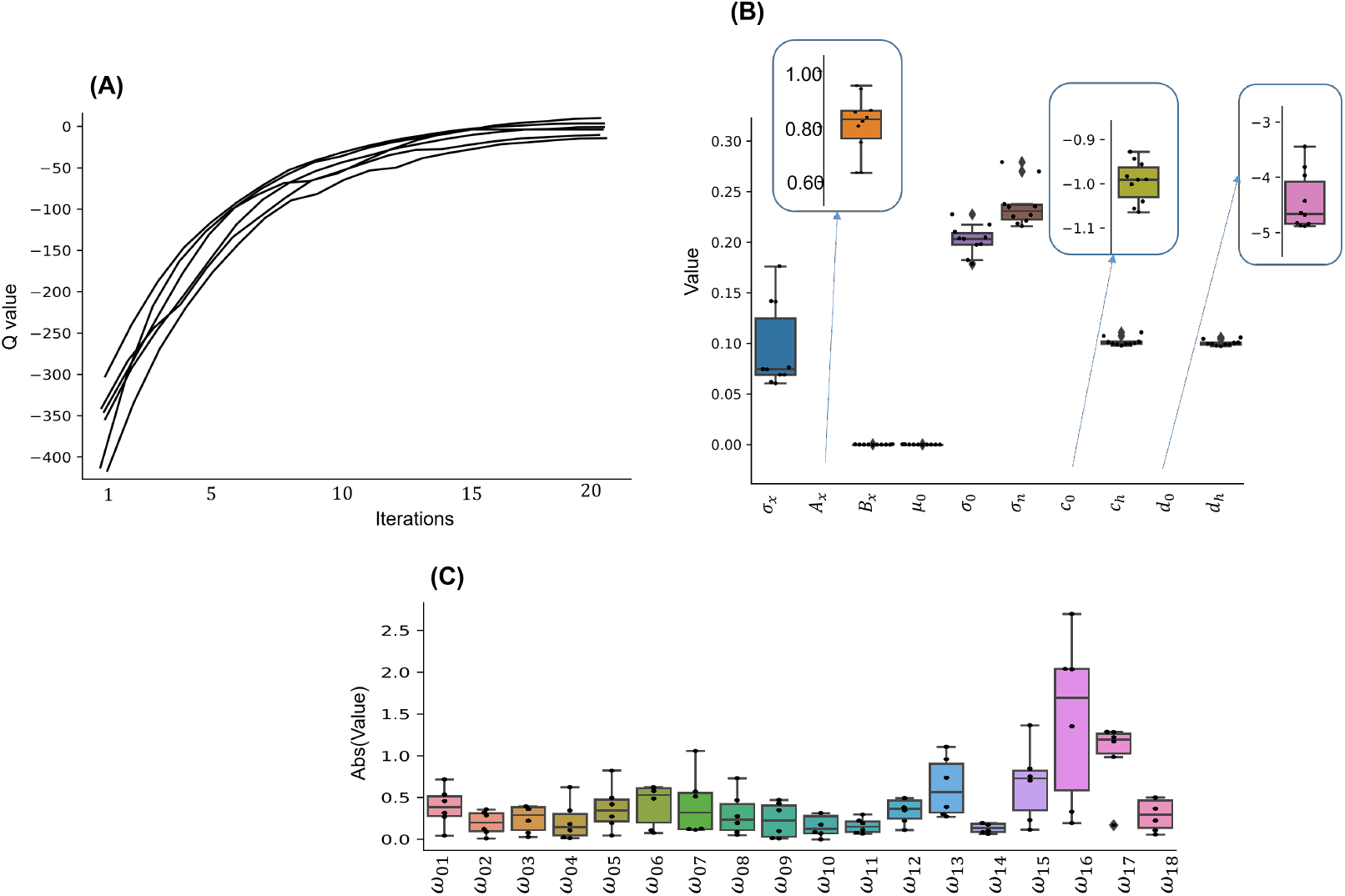
HI-DGD model identification result for experimental data. A) Q values for iterations of the EM algorithm for all the patients. B) Generative processes optimized model parameters for all the patients. C) Discriminative processes optimized model parameters for all the patients.

The optimized model parameters for the neural- and behavioral signals are shown in Figure3.C and Figure3.B, respectively. In the HI-DGD, we assumed that the discriminative process model, *g*(***s**_k_*, ***h***_*k*_; **Ω**), is a linear function of the neural features with additive normal noise with variance *σ_n_* Yousefi et al. (2019). The generative process model, *l*(*x_k_*; **Γ**), is a marked-point process in which its marginal conditional intensity function (CIF) is weighted by the cognitive state, by coefficient *c_x_*, and the history of response times, by coefficient *c_h_*, with a baseline intensity of *c*_0_ Truccolo et al. (2005); Amidi et al. (2019). Similarly, the generation of the associated marks to behavioral signals is parameterized by the cognitive state, by coefficient *d_x_*, and the history of response time, by coefficient *d_h_*, with a baseline of *d*_0_. Notice that we fixed the *c_x_* = 1 and *d_x_* = 1 in the optimization to avoid computational complexity Rezaei et al. (2022a). The variables {*μ*_0_, *σ*_0_} represents the prior knowledge model of the cognitive state with a linear transition process and additive Gaussian noise with variance *σ*_0_. The parameter *σ_x_* is significantly smaller than *σ_n_* which indicates that the cognitive state dynamic is smoother than the actual observed neural activity.

### 3.3. Decoding cognitive state from single trial observations

Here we tested the ability of the HI-DGD ability in decoding the conflict state from single trials of neural recordings (STN-mPFC). Using optimized model parameters, we applied the HI-DGD model to the first 500 msec (i.e., the optimal kernel width for neural features) of neural activities (after the cue time) and inferred the cognitive state associated with each participant. We then passed the mean and variance of decoded cognitive state to a logistic regression model to distinguish between conflict and non-conflict trials, Figure3.B. The logistic regression model can simply draw a border (a threshold) to separate between conflict and non-conflict trials based on the moments of the decoded cognitive state. The time interval used to decode cognitive state is shown by a green shaded area, the first 500ms of neural features, see Figure4.A. The average of the inferred cognitive state is shown in Figure4.B. There is a difference between the distributions of the means in different trial types. This means the mean of the decoded cognitive state for the first 500 msec represents information about the trial types. To evaluate the performance of the decoded cognitive state by HI-DGD in distinguishing between conflict and non-conflict trials, we measured accuracy and f1-score on the prediction result by the logistic regressor for each patient in Figure4.C. The mean accuracy and f1 score for HI-DGD are 77.4% ± 3.0 and 77.5% ± 3.1; respectively. This shows that the cognitive state decoded by HI-DGD reliably and accurately represents conflict processing in the STN in PD patients.

### 3.4. Performance comparison with traditional technique

To identify to what extent the HI-DGD model improves conflict state prediction compared to the traditional analysis, we compared the decoding performance of the HI-DGD model with traditional methods in the literature. One should note that the power of the STN theta band was considered as the main indicator of conflict versus non-conflict choices Ghahremani et al. (2018). Therefore, we compared the HI-DGD model with a baseline model that only uses this feature to distinguish between conflict and non-conflict trials. We used the classifier that was used in the previous section with the STN theta band power as the input feature (consistent with Ghahremani et al. (2018); Zeng et al. (2021)). Similar to the HI-DGD model, we only used the first 500 ms of the neural features after cue time. The accuracy and f1-score of the baseline and the HI-DGD model are shown in Figure4.D. The HI-DGD model significantly improved the prediction of conflict vs. non-conflict trials compared to the baseline model for all the patients(KS-2 Hudson et al. (1992) with p-value 0.002). We also compared the HI-DGD result with the Encoder-Decoder model suggested in Yousefi et al. (2019) in Figure4.D. The plot shows the HI-DGD increased the accuracy and f1-score of the predictions by approximately 5%.

### 3.5. Significant neural features underlying conflict and non-conflict processing

To determine the importance of individual neural features in conflict processing by HI-DGD, we masked all neural features except the feature of interest and used the neural decoder to measure the accuracy of the prediction. In this way, we could identify the contribution of each neural feature in decoding the cognitive state. As shown in Figure4.A, the most significant features are

1. The power of the STN and mPFC channels in the 2-8 Hz frequency band.
2. Power of the mPFC channel in the gamma frequency band (65-95 Hz frequency band).
3. Phase coherency between STN and mPFC in the theta frequency band (2-8 Hz frequency band).
4. Phase coherency between STN and mPFC in the gamma frequency band (65-95 Hz frequency band).

Low-frequency features (2-8 Hz frequency band) have already been noticed in the literature as features that change significantly between conflict and non-conflict trials in the Stroop task Ghahremani et al. (2018); Zeng et al. (2021).

Furthermore, we identified which neural features are necessary for accurate decoding of cognitive states, therefore we tested the performance of the HI-DGD with all neural features against that using only the significant ones identified in Figure6.A. We use the receiver operator characteristic (ROC) curve as an evaluation metric for the HI-DGD performance. The ROC curve is a probabilistic curve that indicates the true prediction rate against the false prediction rate at various threshold values for a classifier in identifying conflict and non-conflict trials. We plotted the ROC curve for HI-DGD performance with all neural features and using only the significant ones in Figure 6.B-C; respectively. The area under the curve (AUC) of the ROC plot is the measure of the ability of the classifier to identify trial types and is used as a summary of the ROC curve. The dashed line in Figure6.B-C, associated with AUC=0.5, indicates the performance of a non-expert classifier that is unable to distinguish between trial types. Therefore, The higher the AUC, the better performance of the model at distinguishing between the trial types. The AUC for HI-DGD performance for the patients is shown in Figure6.D. The non-significant difference between the AUC of HI-DGD with the significant neural features and all neural features shows that the identified significant neural features represent a similar result as the HI-DGD model does with all neural features. We also verified the findings by measuring accuracy and f1 scores for the HI-DGD in distinguishing between the trial types in Figure6. The difference between these measures is negligible.

## 4. Discussion

Both mPFC and STN play crucial roles in conflict processing Zeng et al. (2021). Previous studies reported elevated theta power in STN and mPFC, and increased phase synchrony in the theta band between STN and mPFC in conflict trials Ghahremani et al. (2018); Zeng et al. (2021). A recent study showed increased theta-gamma phase-amplitude coupling (PAC) at STN in conflict trials Zeng et al. (2021). Despite the importance of these features in conflict processing (calculated by group-averaging methods across conflict and non-conflict trials), none of them represent common dynamics of conflict processing at every single trial of observation. Such dynamics, referred to as a cognitive state, represent a low-dimensional representation of the mPFC-STN interaction and link it to the behavioral signal, i.e., the response time. In this study, we developed a Bayesian-based method, namely, heterogeneous input discriminative-generative decoder (HI-DGD) model to infer a cognitive state underlying decision-making from simultaneously recorded EEG from the mPFC and local field potentials from the STN recorded using deep brain stimulation electrodes in PD patients performing a Stroop task. First, we used extensive synthetic data to assess the performance of the model. We showed that the HI-DGD model can infer cognitive states accurately and tracks the dynamics of the data. Second, by applying the HI-DGD model to the experimental data, we showed that the HI-DGD model can predict conflict vs. non-conflict choices using only 500 msec of STN-mPFC recordings. Moreover, the performance of the HI-DGD model in distinguishing between conflict and non-conflict choices was significantly better that using traditional methods. Third, we identified significant neural features estimated by the HI-DGD model: (a) theta band power in both STN and mPFC, (b) theta phase synchrony between mPFC and STN, and (c) gamma power in mPFC. These features match well with those reported in previous studies, indicating the interpretability of the HI-DGD model. We anticipate that the HI-DGD model can be used in closed-loop cognitive systems.

### 4.1. Inferred cognitive state reveals significant neural features underlying conflict and non-conflict choices

Given extensive analysis from previous section, we identified which neural features have significant contributions in distinguishing between conflict and non-conflict choices. As shown in Figure 6.A, the power of the theta frequency band (2-8 Hz) in both STN and mPFC, as well as the phase coherency in the theta band, significantly contributed to the decoding of conflict vs. non-conflict choices. These results of the HI-DGD model are in agreement with those reported in the literature Ghahremani et al. (2018); Zeng et al. (2021), confirming that the inferred cognitive state represents a low dimensional representation of the cognitive state underlying conflict processing in the STN-mPFC neuronal circuit.

Moreover, the HI-DGD model revealed that the power of the gamma band in mPFC is a significant contributor to decoding conflict and non-conflict trials. Gamma activities are mostly associated with contralateral movement such as hand movements Kondylis et al. (2016), or speech production Bush et al. (2022). Speech production, in a verbal Stroop task, is an inevitable part of the task and its onset is marked as the response time. As reported in Ghahremani et al. (2018), the reaction times were significantly longer for conflict trials than non-conflict trials, and gamma activity was locked to the reaction time (speech onset). Therefore, the contribution of gamma activity may originate from speech production during the task.

### 4.2. The HI-DGD enables trial-based detection of conflict and non-conflict states

One of the main advantages of the HI-DGD model, compared to the traditional group-averaging analysis, is that the cognitive state can be inferred (and thus decoded) from individual trials. The model not only enables studying the trial-to-trial variability of cognitive states underlying conflict processing but also provides an opportunity to track the dynamics of these states and design model-based closed-loop cognitive controllers. In Section 3.3, we applied a (trained) neural decoder in the first 500 msec of observed neural activities after the cue time. An interval of 500 msec is sufficient to track at least one cycle of each frequency band, specifically for the theta band power which is a prominent neural feature of conflict processing. We showed in Figure 5 that the HI-DGD model can capture dynamics underlying conflict processing and distinguishes between conflict and non-conflict choices with the mean accuracy of 77.4% (compared to 61.0% using traditional methods). We anticipate that the outcomes of the HI-DGD model can be used to develop closed-loop cognitive controllers in humans.

**Figure 5:**
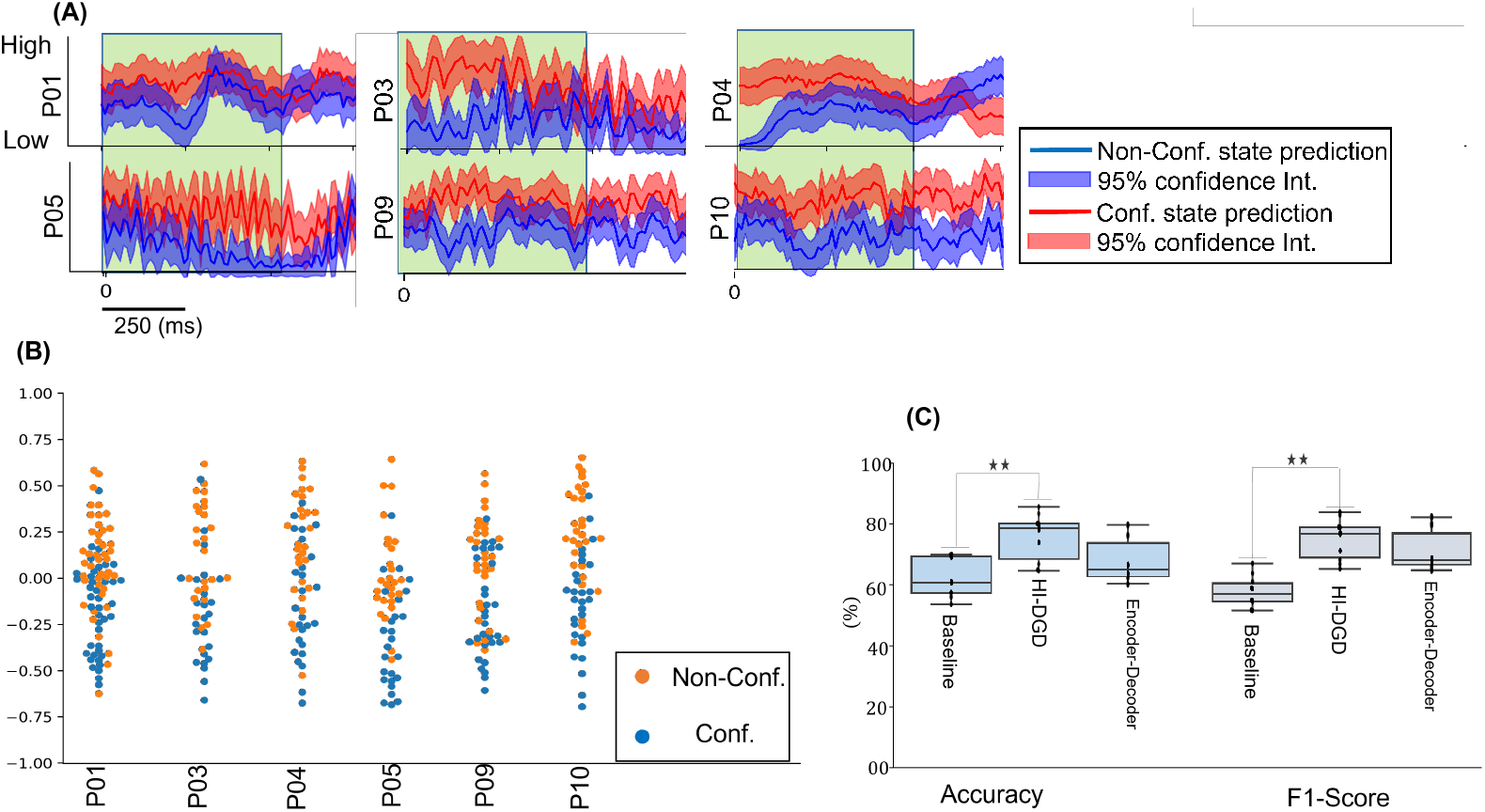
HI-DGD result on the experimental data. A) inferred cognitive stat by HI-DGD. The inferred cognitive state for different trial types of the patients are identified by red (conflict trials) and blue (non-conflict trials). The transparency area identifies the first 500ms of the inferred cognitive state that used for identifying the trial types. B) Mean of the inferred cognitive state for the first 500ms after the cue time for different trial types and each patient. C) Accuracy and F1-score for HI-DGD and the baseline model evaluated on all patients. Hi-DGD outperformed the baseline in both accuracy and F1-score performance measures significantly (p-value 0.002).

**Figure 6:**
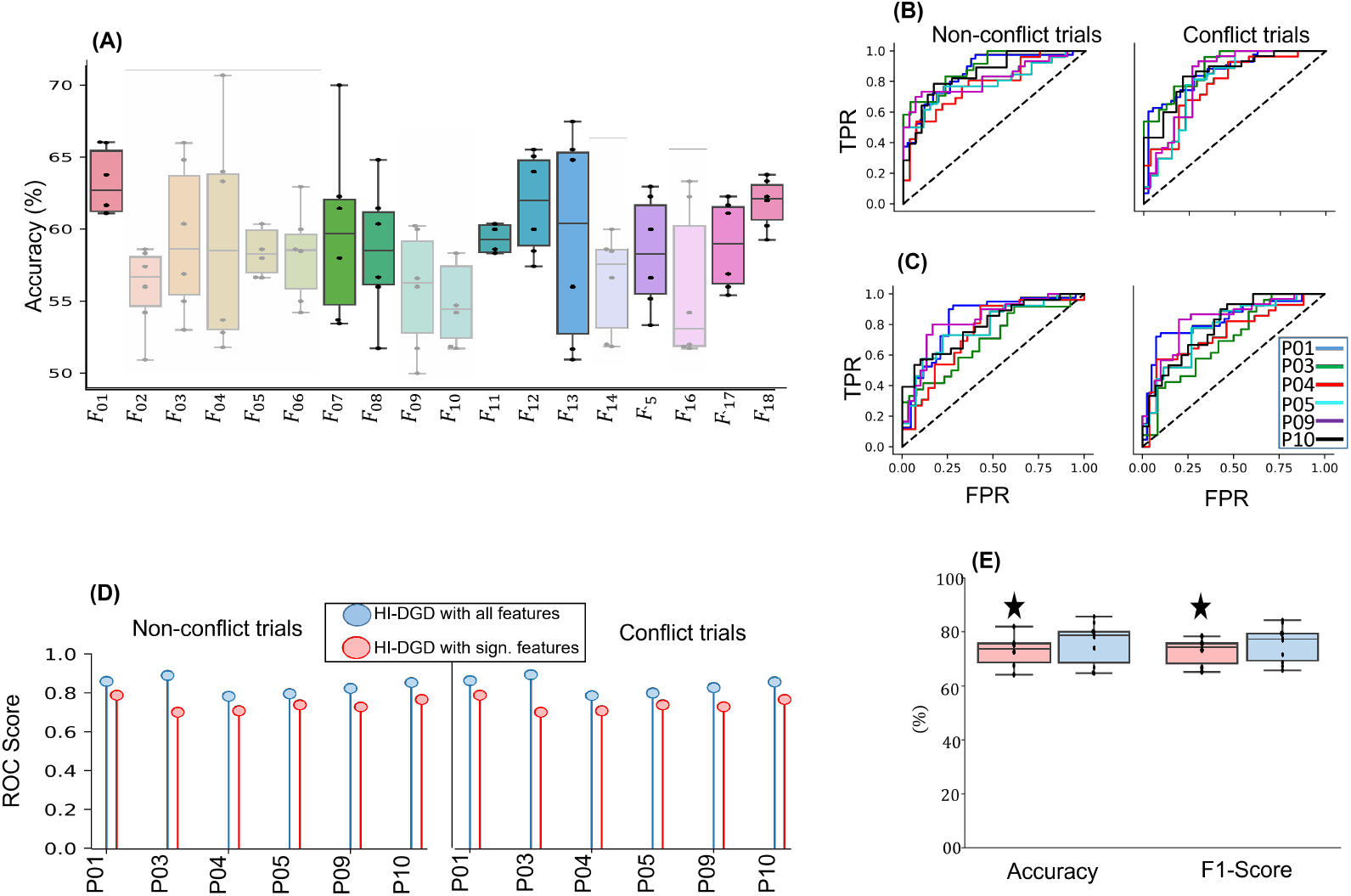
HI-DGD model interpretation and explanation for the experimental data. A) the significant result for each neural feature and all patients. B) The ROC curve for the conflict state prediction with all neural features. C) The ROC curve for the conflict state prediction with the significant neural features highlighted in A). D) ROC score comparison of HI-DGD model with significant (with star) and all neural features evaluated on all patients. E) Accuracy and F1-score for HI-DGD model with significant (with star) and all neural features evaluated on all patients. The accuracy and F1-score performance measures are approximately equal for the HI-DGD model with significant and all neural features.

### 4.3. The HI-DGD outperforms traditional methods in conflict processing

We compared the performance of the HI-DGD model with that using a baseline analysis technique. For the latter, we used a logistic regressor with an input calculated by the power of the theta band of the STN averaged over all trials and across participants (consistent with Ghahremani et al. (2018); Zeng et al. (2021)). Similar to the HI-DGD model, we only used the first 500 ms of observed neural recordings after the cue time. We showed that the HI-DGD model significantly improved the accuracy of the predictions compared to the baseline technique (statistics for accuracy and F1-score, with p-value=0.002).

In developing our modeling framework, the HI-DGD, we hypothesized that there is a low-dimensional dynamical manifold – or, a state process – present in the data which helps to better characterize and potentially understand neural mechanisms of complex cognitive processes. The state process in HI-DGD is directly connected to both generative and discriminative processes and can update its value by either one or two observation(s) at the same time. The discriminative part can therefore be used for decoding a latent state from high-dimensional observed neural recordings Smith and Gales (2002); Burkhart et al. (2020). By considering the discriminative process, the HI-DGD does not need to perform intermediate processing steps such as spike sorting or independent component analysis, which are common practices in preparing neural data Wold et al. (1987). As the HI-DGD uses the discriminative model for high-dimensional data, it offers a more accurate prediction of the cognitive state compared to generative models where the model inadequacy might be problematic Raina et al. (2003); Bernardo et al. (2007). There are many choices to build the discriminative model including deep neural networks (DNNs), convolutional neural networks (CNNs) Albawi et al. (2017), recurrent neural networks (RNNs) Mikolov et al. (2010), and parametric models like the Gaussian process Yu Byron et al. (2009). Unlike high dimensional neural recordings, behavioral signals such as response-time and error rates are usually very low dimensional and have slow temporal dynamics compared to neural signals, thus it is not efficient to use discriminative processes for capturing dynamics of behavioral signals Lasserre et al. (2006); Gordon and Hernández-Lobato (2020). Therefore, we used a generative model, similar to state-space model, for the low dimensional subset of the observed behavioral signals.

## Appendix A. objective function derivation for HI-DGD model identification

We use EM algorithm Dempster et al. (1977) to find maximum likelihood estimates of the model parameters, ***θ*** = {***β*, Ω, *ω***}. The EM algorithm is an established solution to perform maximum likelihood estimation of model parameters when there is an unobservable process or missing observations Yousefi et al. (2019). In the HI-DGD, the state variable ***x_k_*** is the unobserved process, the latent dynamical variable. The EM solution recursively estimates the model parameters ***θ***^(*r*)^ = {***β***^(*r*)^, **Ω**^(*r*)^, **ω**^(*r*)^}, based on an updated posterior distribution of ***x*_0:*K*_** and the observation from the previous EM iteration, ***θ***^(*r*-1)^ = {***β***^(*r*-1)^, **Ω**^(*r*-1)^, **ω**^(*r*-1)^}. The EM includes two steps: expectation and maximization Van Dyk (2000). The expectation step (or *Q* function) is defined by

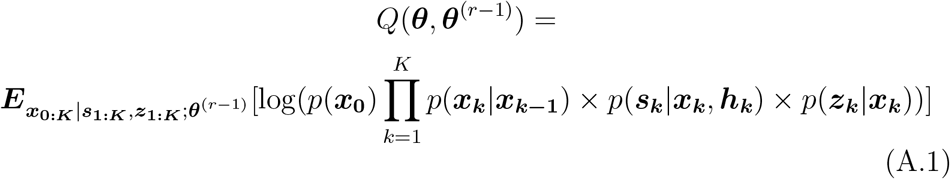

Here after we replace 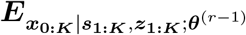 with ***E_K_*** for summarizing the notation. The ***Q*** function can be rewritten as

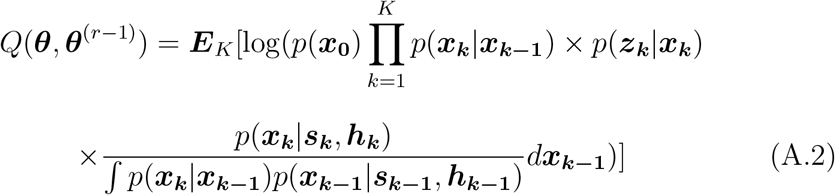

Expanding the *Q* function yields

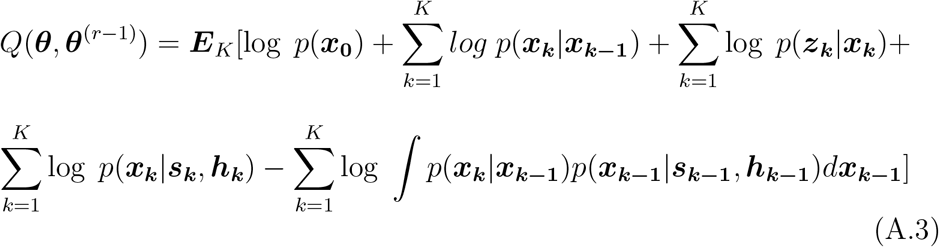

The integration term here can be expressed as

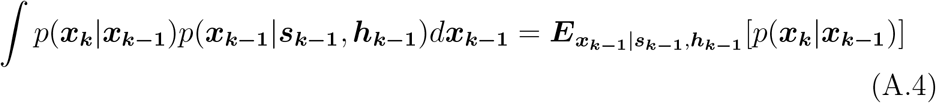

When the state process is linear with an additive Gaussian noise and the prediction process is a multi-variate normal, there is a closed form solution for this expectation. To derive a general solution for nonlinear and non-Gaussian noises, we can rewrite *Q* function as

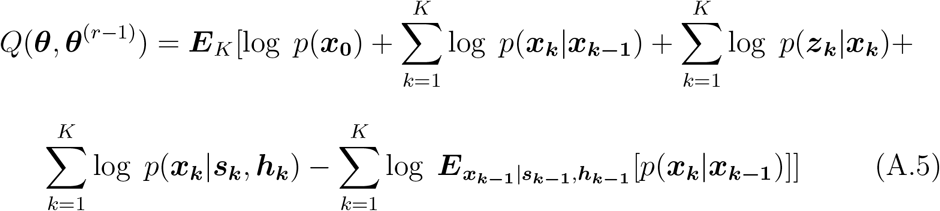

Since log(*E*(*f*(*x*))) ≥ *E*[log(*f*(*x*))], we can exchange the log and expectation operations to yield a lower bound for *Q*, which can be written as

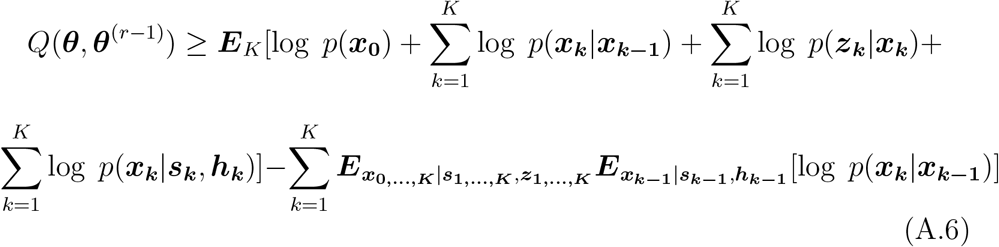

where, the expectation ***x***_*k*-1_ in the last term, defined by the prediction process, is a function of the model free parameters. The second step of the EM is maximization step (M-step) that updates parameter set at iteration (r) by maximizing *Q*. The maximization is defined by

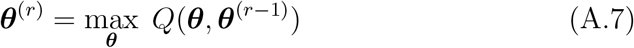

This optimization can be calculated analytically or numerically (e.g. gradient descent Blei et al. (2017)). After each iteration, a new set of parameters are estimated and the EM routine is stopped when a stopping criteria based on the likelihood growth or parameter changes is satisfied.

## Appendix B. simulation data and results

We simulated a group of patients with the assumption that there is a low dimensional dynamics, we referred to it as the cognitive state, that drives both neural activity and behavioral signals. The statistical properties of the cognitive state are different for trial types. Let us assume the cognitive state for trial *j* of *m^th^* patient, *m* = 0,…, *M* is defined by 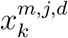, where *d* is the trial type and is modeled by a Bernoulli random variable. Here we selected the Bernoulli parameter 0.5 to make sure we have balanced data from conflict and non-conflict trials. We modeled the state process as a random walk as follows

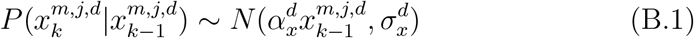

Where 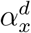 and 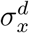 are the transition coefficient and standard deviation for each random walk. We generated two sets of observations. The first one is a high-dimensional and continuous, ***s**_k_*, designed to resemble the dynamics of the LFP/mPFC signals in a cognitive task. Using the state, we then generate a 20-dimensional continuous signal with the conditional distribution defined by

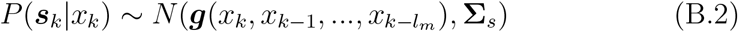

where ***g***(·) is a 20-dimensional vector of non-linear functions, like tanh, and cosine, with an argument defined by a subset of state processes at the current and previous time points. The *l_m_* is the maximum number of data points used in creating *g*(.) function. For each channel of data, we pick a *l* uniformly from 0 to *l_m_* randomly, which is used for data generation. For example, in our simulation, the g_1_ function corresponding to the first element of ***g***, is a tanh(.) with the argument to be +0.8*x*_*k*-1_ +0.6\**x*_*k*-2_+0.4\**x*_k-3_+0.2\**x*_*k*-4_. **Σ**_*s*_ is the stationary covariance matrix with sizer 20 × 20. **Σ**_*s*_’s non-diagonal terms are non-zeros, implying observations across channels are correlated. In our simulation, we picked *a* = .9, *b* = 0, and *σ_x_* = 0.1.

The second observation, *r_k_*, is a low dimensional, discrete, and sparse signal that resembles the dynamics of a behavioral signal in a cognitive task such as response time. *r_k_* can be seen as the time of the agent response to a task and it can be modeled as:

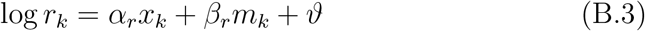

where *α_r_* = 0.9 represent the weight of current state and *β_r_* = 0.1 represent the weight of the error signal *m_k_* in generating log *r_t_. ϑ* is an additive Gaussian noise with a standard deviation of 0.3. We assume this relationship between *r_k_*, conflict state, and error rate because empirically response times typically decline rapidly in non-conflict trials and later tend to reach a lower bound based on subject-specific physiological constraints.

We generate the data for 1000 data points and 5 simulated patients. FigureB.7.A shows a sample of simulated observed data. Our simulation data has a complex observation along with a low-dimensional state process; this setting was purposefully picked to justify the HI-DGD prediction power and build a clear comparison across models. The visualized of the HI-DGD result on the simulated data in FigureB.7.B-E verifies the ability of the HI-DGD model in inferring a latent state from a group of noisy observations.

## Appendix C. Supplementary Figuers

**Figure B.7:**
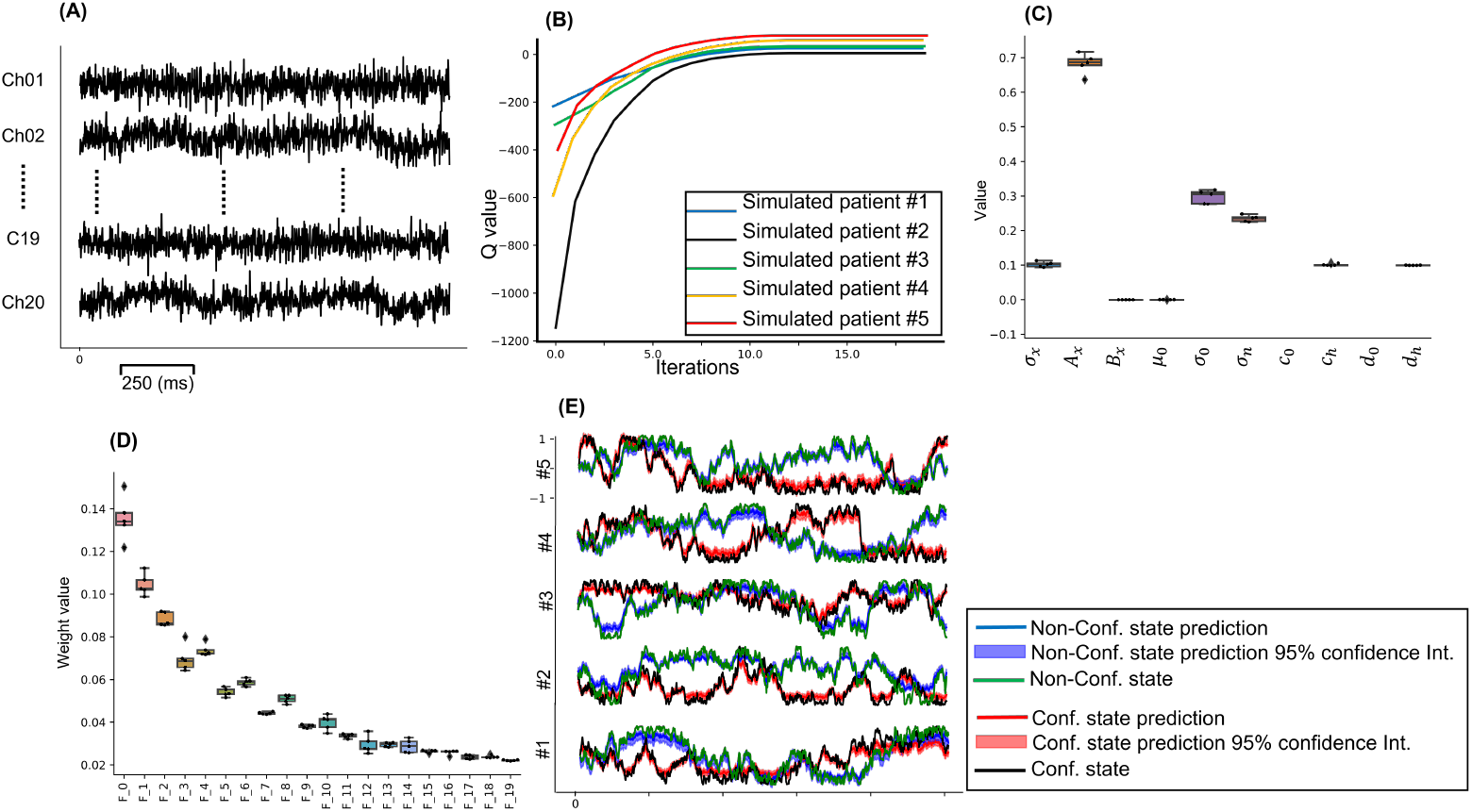
The HI-DGD result on simulation data. A) A sample of simulated neural activity with 20 channels. B) Q values for iterations of the EM algorithm for all the simulated patients. C) Generative processes optimized model parameters for all the patients. D) Discriminative processes optimized model parameters for all the simulated patients. E) inferred cognitive stat by HI-DGD. The inferred hidden state for different trial types of the simulation is identified by red (conflict trials) and blue (non-conflict trials).

**Figure C.8:**
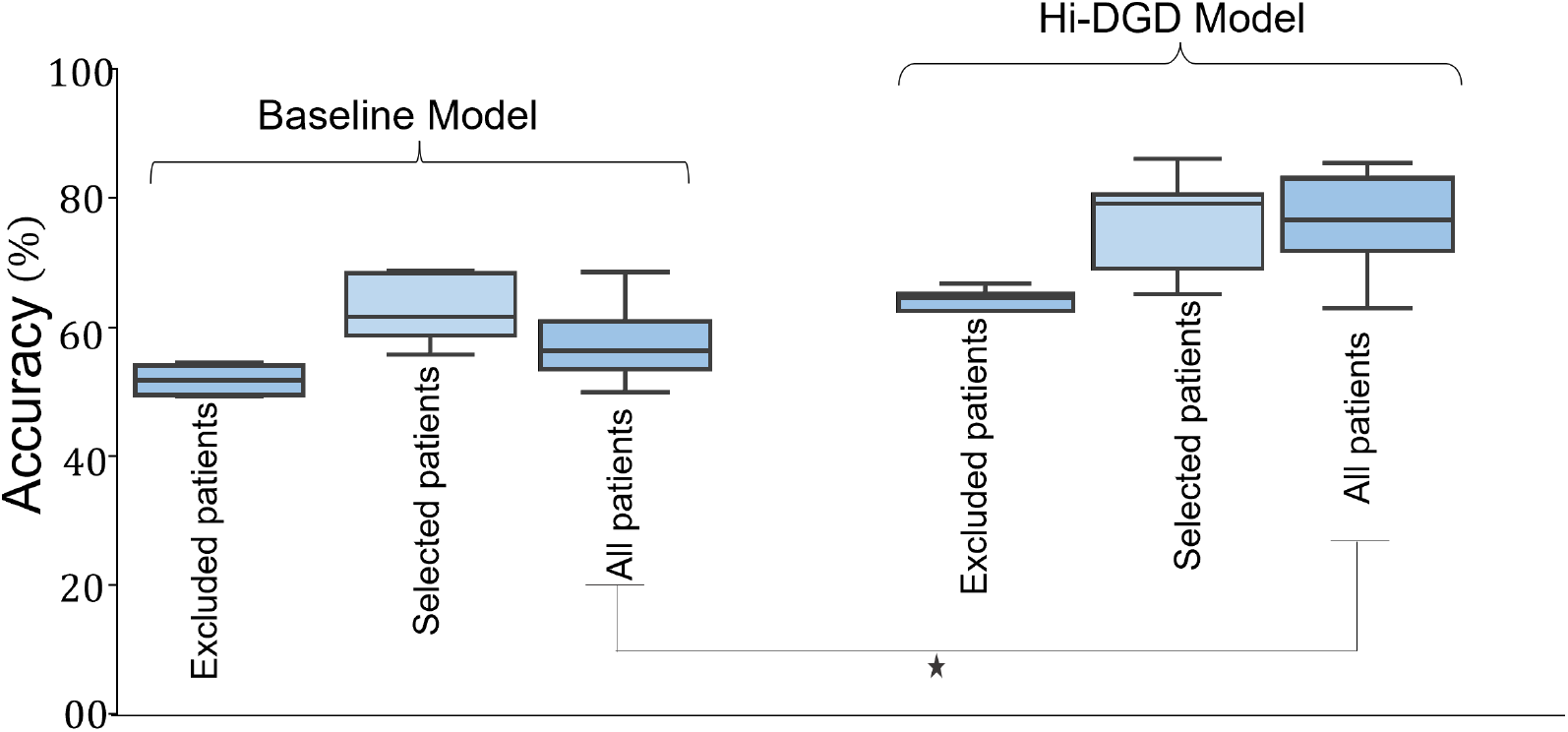
HI-DGD accuracy measure for 10 patients in the experimental data. Accuracy for HI-DGD and the baseline model evaluated on all patients. The accuracy of the proposed HI-DGD is significantly higher than that of the baseline model (p-value ≤ 0.001).

## Notes

### Competing Interest Statement

The authors have declared no competing interest.

## References

Abellán-Nebot, J., Liu, J., Romero, F., 2009. Limitations of the current state space modelling approach in multistage machining processes due to operation variations, in: AIP Conference Proceedings, American Institute of Physics. pp. 231–243.

Albawi, S., Mohammed, T.A., Al-Zawi, S., 2017. Understanding of a convolutional neural network, in: 2017 international conference on engineering and technology (ICET), Ieee. pp. 1–6.

Amidi, Y., Paulk, A.C., Dougherty, D.D., Cash, S.S., Widge, A.S., Eden, U.T., Yousefi, A., 2019. Continuous prediction of cognitive state using a marked-point process modeling framework, in: 2019 41st Annual International Conference of the IEEE Engineering in Medicine and Biology Society (EMBC), IEEE. pp. 2933–2938.

Aron, A.R., Herz, D.M., Brown, P., Forstmann, B.U., Zaghloul, K., 2016. Frontosubthalamic circuits for control of action and cognition. Journal of Neuroscience 36, 11489–11495.

Basu, I., Yousefi, A., Crocker, B., Zelmann, R., Paulk, A.C., Peled, N., Ellard, K.K., Weisholtz, D.S., Cosgrove, G.R., Deckersbach, T., et al., 2021. Closed-loop enhancement and neural decoding of cognitive control in humans. Nature Biomedical Engineering, 1–13.

Bernardo, J., Bayarri, M., Berger, J., Dawid, A., Heckerman, D., Smith, A., West, M., 2007. Generative or discriminative? getting the best of both worlds. Bayesian statistics 8, 3–24.

Blei, D.M., Kucukelbir, A., McAuliffe, J.D., 2017. Variational inference: A review for statisticians. Journal of the American statistical Association 112, 859–877.

Bočková, M., Chládek, J., Jurák, P., Halámek, J., Baláž, M., Rektor, I., 2011. Involvement of the subthalamic nucleus and globus pallidus internus in attention. Journal of neural transmission 118, 1235–1245.

Bočková, M., Lamoš, M., Klimeš, P., Jurák, P., Halámek, J., Goldemundová, S., Baláž, M., Rektor, I., 2020. Suboptimal response to stn-dbs in parkinson’s disease can be identified via reaction times in a motor cognitive paradigm. Journal of Neural Transmission 127, 1579–1588.

Bonnevie, T., Zaghloul, K.A., 2019. The subthalamic nucleus: unravelling new roles and mechanisms in the control of action. The Neuroscientist 25, 48–64.

Brittain, J.S., Watkins, K.E., Joundi, R.A., Ray, N.J., Holland, P., Green, A.L., Aziz, T.Z., Jenkinson, N., 2012. A role for the subthalamic nucleus in response inhibition during conflict. Journal of Neuroscience 32, 13396–13401.

Burkhart, M.C., Brandman, D.M., Franco, B., Hochberg, L.R., Harrison, M.T., 2020. The discriminative kalman filter for bayesian filtering with nonlinear and nongaussian observation models. Neural computation 32, 969–1017.

Bush, A., Chrabaszcz, A., Peterson, V., Saravanan, V., Dastolfo-Hromack, C., Lipski, W.J., Richardson, R.M., 2022. Differentiation of speech-induced artifacts from physiological high gamma activity in intracranial recordings. NeuroImage 250, 118962.

Bush, G., Shin, L.M., 2006. The multi-source interference task: an fmri task that reliably activates the cingulo-frontal-parietal cognitive/attention network. Nature protocols 1, 308–313.

Coleman, T.P., Yanike, M., Suzuki, W.A., Brown, E.N., 2011. A mixed-filter algorithm for dynamically tracking learning from multiple behavioral and neurophysiological measures. The dynamic brain: An exploration of neuronal variability and its functional significance, 3–28.

Dempster, A.P., Laird, N.M., Rubin, D.B., 1977. Maximum likelihood from incomplete data via the em algorithm. Journal of the Royal Statistical Society: Series B (Methodological) 39, 1–22.

Deng, X., Liu, D.F., Kay, K., Frank, L.M., Eden, U.T., 2015. Clusterless decoding of position from multiunit activity using a marked point process filter. Neural computation 27, 1438–1460.

Drummond, N.M., Chen, R., 2020. Deep brain stimulation and recordings: insights into the contributions of subthalamic nucleus in cognition. Neuroimage 222, 117300.

Frank, M.J., 2006. Hold your horses: a dynamic computational role for the subthalamic nucleus in decision making. Neural networks 19, 1120–1136.

Gerstein, G.L., Mandelbrot, B., 1964. Random walk models for the spike activity of a single neuron. Biophysical journal 4, 41–68.

Ghahremani, A., Aron, A.R., Udupa, K., Saha, U., Reddy, D., Hutchison, W.D., Kalia, S.K., Hodaie, M., Lozano, A.M., Chen, R., 2018. Event-related deep brain stimulation of the subthalamic nucleus affects conflict processing. Annals of neurology 84, 515–526.

Goodwin, G.C., Graebe, S.F., Salgado, M.E., et al., 2001. Control system design. volume 240. Prentice Hall Upper Saddle River.

Gordon, J., Hernández-Lobato, J.M., 2020. Combining deep generative and discriminative models for bayesian semi-supervised learning. Pattern Recognition 100, 107156.

Gramfort, A., Luessi, M., Larson, E., Engemann, D.A., Strohmeier, D., Brodbeck, C., Goj, R., Jas, M., Brooks, T., Parkkonen, L., et al., 2013. Meg and eeg data analysis with mne-python. Frontiers in neuroscience, 267.

Güçlütürk, Y., Güçlü, U., Seeliger, K., Bosch, S., van Lier, R., van Gerven, M.A., 2017. Reconstructing perceived faces from brain activations with deep adversarial neural decoding. Advances in neural information processing systems 30.

Haynes, W.I., Haber, S.N., 2013. The organization of prefrontal-subthalamic inputs in primates provides an anatomical substrate for both functional specificity and integration: implications for basal ganglia models and deep brain stimulation. Journal of Neuroscience 33, 4804–4814.

Hudson, R.R., Boos, D.D., Kaplan, N.L., 1992. A statistical test for detecting geographic subdivision. Molecular biology and evolution 9, 138–151.

Karush, J., 1961. On the chapman-kolmogorov equation. The Annals of Mathematical Statistics 32, 1333–1337.

Kelley, R., Flouty, O., Emmons, E.B., Kim, Y., Kingyon, J., Wessel, J.R., Oya, H., Greenlee, J.D., Narayanan, N.S., 2018. A human prefrontal-subthalamic circuit for cognitive control. Brain 141, 205–216.

Kingma, D.P., Welling, M., 2013. Auto-encoding variational bayes. arXiv preprint arXiv:1312.6114.

Kondylis, E.D., Randazzo, M.J., Alhourani, A., Lipski, W.J., Wozny, T.A., Pandya, Y., Ghuman, A.S., Turner, R.S., Crammond, D.J., Richardson, R.M., 2016. Movement-related dynamics of cortical oscillations in parkinson’s disease and essential tremor. Brain 139, 2211–2223.

Lasserre, J.A., Bishop, C.M., Minka, T.P., 2006. Principled hybrids of generative and discriminative models, in: 2006 IEEE Computer Society Conference on Computer Vision and Pattern Recognition (CVPR’06), IEEE. pp. 87–94.

Lutz, A., Lachaux, J.P., Martinerie, J., Varela, F.J., 2002. Guiding the study of brain dynamics by using first-person data: synchrony patterns correlate with ongoing conscious states during a simple visual task. Proceedings of the national academy of sciences 99, 1586–1591.

Marshall, S.P., 2007. Identifying cognitive state from eye metrics. Aviation, space, and environmental medicine 78, B165–B175.

Mikolov, T., Karafiát, M., Burget, L., Cernocký, J., Khudanpur, S., 2010. Recurrent neural network based language model., in: Interspeech, Makuhari. pp. 1045–1048.

Nolte, G., Bai, O., Wheaton, L., Mari, Z., Vorbach, S., Hallett, M., 2004. Identifying true brain interaction from eeg data using the imaginary part of coherency. Clinical neurophysiology 115, 2292–2307.

Panzeri, S., Macke, J.H., Gross, J., Kayser, C., 2015. Neural population coding: combining insights from microscopic and mass signals. Trends in cognitive sciences 19, 162–172.

Prerau, M.J., Smith, A.C., Eden, U.T., Kubota, Y., Yanike, M., Suzuki, W., Graybiel, A.M., Brown, E.N., 2009. Characterizing learning by simultaneous analysis of continuous and binary measures of performance. Journal of neurophysiology 102, 3060–3072.

Prerau, M.J., Smith, A.C., Eden, U.T., Yanike, M., Suzuki, W.A., Brown, E.N., 2008. A mixed filter algorithm for cognitive state estimation from simultaneously recorded continuous and binary measures of performance. Biological cybernetics 99, 1–14.

Raina, R., Shen, Y., Mccallum, A., Ng, A., 2003. Classification with hybrid generative/discriminative models. Advances in neural information processing systems 16.

Rezaei, M.R., Arai, K., Frank, L.M., Eden, U.T., Yousefi, A., 2021. Real-time point process filter for multidimensional decoding problems using mixture models. Journal of neuroscience methods 348, 109006.

Rezaei, M.R., Gillespie, A.K., Guidera, J.A., Nazari, B., Sadri, S., Frank, L.M., Eden, U.T., Yousefi, A., 2018. A comparison study of point-process filter and deep learning performance in estimating rat position using an ensemble of place cells, in: 2018 40th Annual International Conference of the IEEE Engineering in Medicine and Biology Society (EMBC), IEEE. pp. 4732–4735.

Rezaei, M.R., Hadjinicolaou, A.E., Cash, S.S., Eden, U.T., Yousefi, A., 2022a. Direct discriminative decoder models for analysis of highdimensional dynamical neural data. Neural Computation 34, 1100–1135.

Rezaei, M.R., Popovic, M.R., Lankarany, M., Yousefi, A., 2022b. Deep discriminative direct decoders for high-dimensional time-series analysis. arXiv preprint arXiv:2205.10947.

Robert, M., Faraj, A., McAllister, M.K., Rivot, E., 2010. Bayesian statespace modelling of the de lury depletion model: strengths and limitations of the method, and application to the moroccan octopus fishery. ICES Journal of Marine Science 67, 1272–1290.

Ruff, D.A., Ni, A.M., Cohen, M.R., 2018. Cognition as a window into neuronal population space. Annual review of neuroscience 41, 77.

Sani, O.G., Abbaspourazad, H., Wong, Y.T., Pesaran, B., Shanechi, M.M., 2021. Modeling behaviorally relevant neural dynamics enabled by preferential subspace identification. Nature Neuroscience 24, 140–149.

Sani, O.G., Yang, Y., Lee, M.B., Dawes, H.E., Chang, E.F., Shanechi, M.M., 2018. Mood variations decoded from multi-site intracranial human brain activity. Nature biotechnology 36, 954–961.

Scarpina, F., Tagini, S., 2017. The stroop color and word test. Frontiers in psychology 8, 557.

Smith, N.D., Gales, M.J., 2002. Using svms and discriminative models for speech recognition, in: 2002 IEEE International Conference on Acoustics, Speech, and Signal Processing, IEEE. pp. I–77.

Suzuki, W.A., Brown, E.N., 2005. Behavioral and neurophysiological analyses of dynamic learning processes. Behavioral and cognitive neuroscience reviews 4, 67–95.

Truccolo, W., Eden, U.T., Fellows, M.R., Donoghue, J.P., Brown, E.N., 2005. A point process framework for relating neural spiking activity to spiking history, neural ensemble, and extrinsic covariate effects. Journal of neurophysiology 93, 1074–1089.

Van Der Merwe, R., 2004. Sigma-point Kalman filters for probabilistic inference in dynamic state-space models. Oregon Health & Science University.

Van Dyk, D.A., 2000. Fitting mixed-effects models using efficient em-type algorithms. Journal of Computational and Graphical Statistics 9, 78–98.

Villemure, C., Bushnell, C.M., 2002. Cognitive modulation of pain: how do attention and emotion influence pain processing? Pain 95, 195–199.

Wessel, J.R., Waller, D.A., Greenlee, J.D., 2019. Non-selective inhibition of inappropriate motor-tendencies during response-conflict by a fronto-subthalamic mechanism. Elife 8, e42959.

Wold, S., Esbensen, K., Geladi, P., 1987. Principal component analysis. Chemometrics and intelligent laboratory systems 2, 37–52.

Wood, E., Fellows, M., Donoghue, J., Black, M., 2004. Automatic spike sorting for neural decoding, in: The 26th Annual International Conference of the IEEE Engineering in Medicine and Biology Society, IEEE. pp. 4009–4012.

Wu, F., Zilberstein, S., Jennings, N.R., 2013. Monte-carlo expectation maximization for decentralized pomdps, in: Proceedings of the 23rd international joint conference on artificial intelligence (IJCAI), pp. 397–403.

Yousefi, A., Basu, I., Paulk, A.C., Peled, N., Eskandar, E.N., Dougherty, D.D., Cash, S.S., Widge, A.S., Eden, U.T., 2019. Decoding hidden cognitive states from behavior and physiology using a bayesian approach. Neural computation 31, 1751–1788.

Yu Byron, M., Cunningham John, P., Santhanam Gopal, R.S.I., Shenoy Krishna, V., 2009. Sahani maneesh. gaussian-process factor analysis for low-dimensional single-trial analysis of neural population activity. Journal of Neurophysiology, 614–35.

Zeng, K., Drummond, N.M., Ghahremani, A., Saha, U., Kalia, S.K., Hodaie, M., Lozano, A.M., Aron, A.R., Chen, R., 2021. Fronto-subthalamic phase synchronization and cross-frequency coupling during conflict processing. NeuroImage 238, 118205.

